# The gut microbiome and child mental health: a population-based study

**DOI:** 10.1101/2022.08.15.502771

**Authors:** Robert Kraaij, Isabel K. Schuurmans, Djawad Radjabzadeh, Henning Tiemeier, Timothy G. Dinan, André G. Uitterlinden, Manon Hillegers, Vincent W.V. Jaddoe, Liesbeth Duijts, Henriette Moll, Fernando Rivadeneira, Carolina Medina-Gomez, Pauline W. Jansen, Charlotte A.M. Cecil

**Affiliations:** Department of Internal Medicine, Erasmus MC, University Medical Center, Rotterdam, The Netherlands; Department of Epidemiology, Erasmus MC, University Medical Center, Rotterdam, The Netherlands; The Generation R Study Group, Erasmus MC, University Medical Center, Rotterdam, The Netherlands; Department of Child and Adolescent Psychiatry / Psychology, Erasmus MC, University Medical Center, Rotterdam, The Netherlands; Department of Social and Behavioral Sciences, Harvard. T.H. Chan School of Public Health, Boston, MA, USA; APC Microbiome Ireland, University College Cork, Cork, Ireland; Department of Psychiatry and Neurobehavioral Science, University College Cork, Cork, Ireland; Department of Pediatrics, Erasmus MC, University Medical Center, Rotterdam, The Netherlands; Department of Pediatrics, divisions of Respiratory Medicine and Allergology, and Neonatology, Erasmus MC, University Medical Center Rotterdam, Rotterdam, the Netherlands

## Abstract

The link between the gut microbiome and the brain has gained increasing scientific and public interest for its potential to explain psychiatric risk. While differences in gut microbiome composition have been associated with several mental health problems, evidence to date has been largely based on animal models and human studies in small sample sizes. Here, we aimed to systematically characterize associations of the gut microbiome with overall psychiatric symptoms as well as with specific domains of emotional and behavioral problems, assessed via the maternally rated Child Behavior Checklist in 1,784 ten-year-old children from the multi-ethnic, population-based Generation R Study. While we observe lower gut microbiome diversity in case of more overall and specific mental health problems, associations were not significant. Likewise, we did not identify any taxonomic feature associated with mental health problems after multiple testing correction, although nominally significant findings indicated depletion of genera previously associated with psychiatric disorders, including *Anaerotruncus, Anaeroplasma*, and *Hungatella*. The identified compositional abundance differences were found to be similar across all mental health problems. Finally, we did not find any specific microbial functions that were enriched in relation to children with mental health problems. In conclusion, based on the largest sample examined to date, we found no clear evidence of associations between gut microbiome diversity, taxonomies or functions and mental health problems in the general pediatric population. In future, the use of longitudinal designs with repeated measurements microbiome and psychiatric outcomes will be critical to clarify any emerging associations between the gut microbiome and mental health from early life to adulthood.

## INTRODUCTION

The relationship between the gut microbiome and mental health has drawn immense scientific and public interest over recent years (1), spurred by an increased understanding of the key role microbiome may play in mediating communication between the gut and the brain (the so-called ‘gut-brain’ axis (2)). The gut microbiome influences the brain via multiple pathways, including neurotransmitter synthesis (e.g., serotonin), activation of the immune system, production of neuroactive metabolites (e.g., short-chain fatty acids) and vagus nerve stimulation (3). Furthermore, several environmental factors that influence the brain and psychiatric risk have also been shown to impact gut dysbiosis (i.e., imbalance of the gut microbiome), including stress exposure, medications and diet (2). As such, the gut microbiome has emerged as a promising potential mechanism underlying individual differences in brain function, behavior, and psychiatric risk.

Most studies on the gut-brain axis to date have been based on experimental work in animals, demonstrating the importance of gut bacteria for neurodevelopment and behavior, including learning and memory, social interactions, stress response, and anxiety- and depressive-like behaviors (4). A causal effect of the microbiome on the brain has further been supported by fecal microbial transplantation (FMT) studies, which show that translocation of fecal bacteria from human donors with a psychiatric disorder (e.g., depression, anxiety or schizophrenia) associate with reduced microbial diversity and increased psychiatric symptoms in animals (5). The human literature has been sparser and almost entirely comprised of patient studies. These have focused mainly on autism spectrum disorder, reporting lower abundances of *Enterococcus, Escherichia coli, Bacteroides*, and *Bifidobacterium* in patients compared to healthy controls (6); or on major depressive disorder, showing less abundance of *Prevotellaceae, Coprococcus*, and *Faecalibacterium* (7). While consistent and independently replicated findings for depression have recently begun to emerge from large adult population-based studies (8, 9), other psychiatric symptoms have not yet been consistently examined. Yet, preliminary evidence of gut microbial dysbiosis has been found in relation to disorders such attention-deficit hyperactivity disorder, schizophrenia, and generalized anxiety disorder (for a review of available evidence, see (1) and (10)). Besides associations with clinical disorders in adults, a smaller set of studies in infants have reported associations with subclinical mental health (i.e., emotional and behavioral problems) and temperament (e.g., (11) and (12)).

The current evidence base is limited in four key ways. First, research to date on gut microbiome and mental health has been based on modest sample sizes (i.e., an average number of 42 cases per study (10)), which are subject to limitations such as selection bias, unclear generalizability of findings, lack of adequate control for confounding, and low statistical power to detect associations of small effect. Second, studies have varied widely in methodology, including the use of multiple testing correction, adjustment for covariates, and the analysis of gut microbiome at different taxonomic levels, further limiting comparability of findings. This has led to increased calls to move towards larger, well-designed and better powered studies, examining the gut microbiome at multiple levels (e.g., from global diversity measures to individual taxonomic units) (1). Third, the populations examined were either infants or adults, while psychiatric symptoms typically have their onset in childhood and adolescence as emotional and behavioral problems (13). Therefore, it remains unclear whether associations reported may also be evident in childhood. Finally, studies almost exclusively focused on single psychiatric outcomes, despite evidence that psychiatric symptoms often co-occur. As such, it has not been possible to establish whether reported gut microbiome differences may be unique or shared across psychiatric symptoms.

To address these gaps, we examined cross-sectional associations between the gut microbiome and common mental health problems in a general population cohort of almost 1,800 ten-year old children. We utilize a comprehensive approach to characterize associations between gut microbiome composition at different taxonomic levels (alpha diversity measures, genus level, and functional analyses) and overall psychiatric problems in children. As follow-up analyses, we examined associations of the gut microbiome with eight specific domains of emotional and behavioral symptoms.

## METHODS

### The Generation R Study

The Generation R Study is a population-based prospective multi-ethnic cohort from fetal life onward conducted in the city of Rotterdam (14). The study was designed to identify early environmental and genetic factors and causal pathways underlying growth and development during childhood. The Generation R Study recruited 9,749 pregnant women, of whom more than 70% kept on participating after birth, undergoing several rounds of follow-up. Ethics approval was obtained from the Medical Ethical Committee of Erasmus MC (MEC-2012-165) and written informed consent was obtained from all participants’ parents on behalf of their children. All methods were performed in accordance with the Declaration of Helsinki. In total, 5,862 children participated in the wave at age 10 years, of which 2,526 children returned a stool sample that could be included in the microbiome dataset (15) and of which 5,523 mothers returned a valid Child Behavior Checklist (CBCL) (14). After combining participants with an included stool sample and a valid CBCL, 1,784 participants remained to be analyzed (flowchart in Supplementary Table 1).

**Table 1.** Baseline characteristics of the total cohort participating in the age 10 years data collection wave compared to the sub-cohort with child mental health phenotype and microbiome data.

|  | CBCL at age10 and<br>microbiome | Participated in CBCL at age 10 | *p* |
| --- | --- | --- | --- |
| *N* | 1,784 | 5,523 |  |
| Age, years (mean (SD)) | 9.8 (0.29) | 9.7 (0.33) | <.001 |
| Sex, girls (%) | 48.8 | 50.1 | .342 |
| BMI, kg/m<sup>2</sup> (mean (SD)) | 17.2 (2.43) | 17.4 (2.55) | .012 |
| Antibiotic use (%) | 7.3 |  |  |
| Last month (%) | 1.2 |  |  |
| 1 - 3 months ago (%) | 2.0 |  |  |
| 3 months to 1 year ago (%) | 4.1 |  |  |
| Education-level mother at child birth |  |  | .069 |
| Primary education (%) | 4.4 | 5.0 |  |
| Secondary (%) | 36.1 | 37.5 |  |
| Higher (%) | 54.5 | 51.5 |  |
| Missing (%) | 5.0 | 6.0 |  |
| Country of origin |  |  | .208 |
| Dutch (%) | 66.3 | 64.4 |  |
| Non-Dutch (%) | 32.9 | 34.4 |  |
| Missing (%) | 0.8 | 1.2 |  |

Educational level mother is six categories, primary presents the first two, secondary the second two, and higher the last two. Continuous data was compared using t-statistics, while categorical data was compared using Chi-square statistics. CBCL: Child Behavior Checklist, the child mental health phenotype; SD: standard deviation.

### Gut microbiome

Stool samples were collected at a mean age of 9.8 years (SD = 0.3). DNA isolation and 16S rRNA sequencing were performed as described (15). Amplicon sequence variants (ASVs) were derived from the sequencing data using the DADA2 profiling pipeline (version 1.18.0) (16). Full description of the profiling pipeline is provided in the supplementary information. In brief, the V3 and V4 variable regions of the 16S rRNA gene were amplified and sequenced, barcodes and primers were removed from the sequence reads and ASV clustering and denoising was performed. Remaining chimeric sequences were removed and taxonomies were assigned to the ASVs using the RDP naive Bayesian classifier (17), which was trained on the SILVA version 138.1 microbial database (18). Initial ASV filtering was performed based on abundance (0.05% of the total reads) and prevalence (present in at least 1% of the samples). An imputed functional metagenome table was obtained using the PICRUSt 2.0 tool (19). A species and genus-level phyloseq objects were obtained by aggregation of the phyloseq object based on the ASV-level taxonomy table.

### Child mental health problems

Child mental health problems were assessed using the maternally rated Child Behavior Checklist 6-18 (CBCL/6-18) at age 10. Mothers rated various emotional (i.e., internalizing) and behavioral (i.e., externalizing) problems of the child in the previous six months using 118 questions on a three-point scale (0 = not true, 1 = somewhat true, 2 = very true). Based on these items, two broadband scales (internalizing and externalizing problems) and a total problem scale were derived, as well as eight empirical syndrome scales (i.e., anxious/depressed, withdrawn/depressed, somatic complaints, social problems, thought problems, attention problems, rule-breaking behavior and aggressive behavior) (20).

### Other variables

Age was recorded as the date of stool sample production. Sex was obtained through midwifes at birth (21). Body mass index (BMI) was calculated from weight and length recorded during a research visit when children were 10 years old. Antibiotics use was measured at the time of stool collection using self-report, and was categorized into three categories (i.e., last month, 1 to 3 months ago, 3 months to a year age). Education level of the mother at birth was assessed by a questionnaire (21) and defined by the highest attained educational level and classified into six categories (i.e., no education; primary education; lower vocational training or intermediate general school; >3 years secondary school or intermediate vocational training or higher vocational training; bachelor’s degree; higher academic education or PhD). Five technical covariates were included in all analyses, being i) the number of days a stool sample had been in the regular mail during sample collection, ii) season of stool sample production (winter, spring, summer, fall), iii) one of two DNA isolation batches that were observed during dataset generation, iv) one of three sequencing run batches, and v) the number of sequencing reads. Genetic principal components (PCs) were derived from DNA, in order to adjust for population stratification. Finally, country of origin of the participant was assessed based on self-reported country of birth of four grandparents (21). A participant was deemed to be of non-Dutch origin if one parent was born abroad. If both parents were born abroad, the country of birth of the participant’s mother defined the ethnic background.

### Analyses

Analyses were performed in R version 4.1.0 (22) with Bioconductor 3.13 (23) adjusting for age, sex, BMI, self-reported antibiotics use, maternal education, technical covariates, and first 10 genetic PCs as covariates (see Table 1 for details). The only covariate with missing data (maternal education) was imputed by multiple imputation using chained equations with the “mice” package in R (24). 30 datasets were generated using 100 iterations. Pooled estimates were obtained using Rubin’s rules (25), and if pooling functions were not available, median statistics were reported. As microbiome data is compositional and zero-inflated, zero abundances were imputed using the zCompositions package (26) and transformed by centered log ratio transformation. Child mental health problems were square root transformed to approach normality.

Analyses were performed using a hierarchical approach, first examining overall child mental health problems, and subsequently specific child mental health problems. As such, for our primary aim, we assessed associations with broadband (i.e., internalizing and externalizing) and total problems scales; and as follow-up analyses, we examined associations with empirical syndrome scales (i.e., anxious/depressed, withdrawn/depressed, somatic complaints, social problems, thought problems, attention problems, rule-breaking behavior and aggressive behavior). Analyses proceeded in three steps focusing on different taxonomic levels, as described below.

#### Step 1. Associations with alpha diversity indices

Associations of alpha diversities (i.e., richness (27), Shannon diversity (28), and inverse Simpson index (29)) with square root transformed child mental health problems were examined using linear regression models. Alpha diversity indices are one-value summaries of a sample’s microbiome profile that reflect the number of observed species and their mutual distribution.

#### Step 2. Associations with gut microbiome profiles

Univariate (i.e., per taxon) differential abundance analyses and multivariate (i.e., including all taxa in the profile) analyses were performed on the genus-level abundance table. Abundance data were transformed to centered log ratios after imputing zero read abundances to reduce biases introduced by the compositional nature of the microbiome data. Univariate associations of single taxa with child mental health phenotypes were determined by ANCOM-BC differential abundance analysis (30) using the square root transformed child mental health phenotypes. Univariate taxon analyses determine the associations with single taxa without considering the entire microbiome profile and mutual relations between taxa. Multivariate composition was associated with child mental health problems by PERMANOVA (“adonis” function in R library vegan (31)) using the pairwise Euclidean distance matrix of the centered log ratio profiles (beta diversity) and adding the square root transformed phenotype as last variable into the base model. PERMANOVA estimates whether the microbiome profiles of samples within a category are more similar (cluster together) than between categories using the entire profile to determine similarities between samples.

#### Step 3. Functional pathways

MetaCyc relative functional pathway abundances (32) were associated using the base model and linear regression with the square root of child mental health phenotypes as outcomes. Based on the taxa present in each microbiome profile, PICRUSt predicts the level of enzymes that are present in each sample using databases, MetaCyc pathway abundances are then derived from these predicted enzyme levels (32).

#### Multiple testing

Multiple testing was controlled for by Benjamini-Hochberg correction (33) within each analysis, which were run separately in each taxonomic level for (i) primary analyses (i.e., 3 overall scales), and (ii) follow-up analyses (i.e., 8 specific scales). Results were considered significant for q (corrected *p*-value) < 0.05 and nominally significant for *p* < 0.01 and q > 0.05.

## RESULTS

Sample characteristics are displayed in Table 1. Mean age of the participants was 9.8 years with a narrow range (SD = 0.3 years) and mean BMI was 17.2 kg/m^2^. A total of 16 ethnicities were recorded with Dutch (66.3%), European (7.5%), Turkish (3.7%) and Moroccan (3.4%) being the most common. Microbiome characteristics are displayed in Supplementary Table 2. A total of 305 different species-level ASVs were detected in the dataset after filtering. Each sample contained on average 84.2 different species-level ASVs. At genus-level, 188 different genera were detected; while on average 61.1 genera were present in each sample.

**Table 2.** Associations between child mental health phenotypes and overall gut microbiome composition (genus-level)

|  | Microbiome richness |  |  | Shannon Diversity |  |  | inversed Simpson diversity |  |  |
| --- | --- | --- | --- | --- | --- | --- | --- | --- | --- |
|  | B | se | p | B | se | p | B | se | p |
| Broadband and total scales |  |  |  |  |  |  |  |  |  |
| Internalizing problems | -0.941 | 0.665 | .157 | -0.012 | 0.008 | .143 | -0.530 | 0.288 | .066 |
| Externalizing problems | 0.185 | 0.636 | .771 | 0.001 | 0.008 | .884 | 0.115 | 0.276 | .676 |
| Total problems | -0.182 | 0.441 | .680 | -0.005 | 0.006 | .369 | -0.177 | 0.191 | .355 |
| Broadband and total scales |  |  |  |  |  |  |  |  |  |
| Anxious/depressed | -1.361 | 0.803 | .091 | -0.015 | 0.010 | .134 | -0.668 | 0.349 | .055 |
| Withdrawn/depressed | -0.053 | 0.962 | .956 | -0.005 | 0.012 | .696 | -0.062 | 0.418 | .881 |
| Somatic complaints | -0.525 | 0.923 | .570 | -0.007 | 0.012 | .532 | -0.418 | 0.401 | .297 |
| Social problems | -0.276 | 0.872 | .752 | -0.014 | 0.011 | .208 | -0.513 | 0.379 | .176 |
| Thought problems | -0.107 | 0.888 | .904 | -0.009 | 0.011 | .439 | -0.369 | 0.385 | .338 |
| Attention problems | 0.495 | 0.776 | .523 | 0.004 | 0.010 | .706 | 0.261 | 0.337 | .438 |
| Rule-breaking behavior | 0.211 | 1.043 | .840 | 0.001 | 0.013 | .915 | 0.047 | 0.453 | .918 |
| Aggressive behavior | 0.116 | 0.694 | .867 | 0.001 | 0.009 | .872 | 0.133 | 0.301 | .658 |

Linear regression of child mental health phenotypes and gut microbiome alpha diversities. Model: alpha diversity measure ~ sqrt(child mental health phenotype) + age + sex + BMI + self-reported antibiotics use + maternal education + time in mail + season of stool production + DNA isolation batch + sequencing run batch + number of sequencing reads + first 10 genetic PCs. Values are pooled from 30 imputed datasets. Alpha: alpha diversity metric; sqrt: square root transformed; PCs: principal components; se: standard error.

### Child mental health and gut microbiome diversities

After adjusting for age, sex, BMI, self-reported antibiotics use, maternal education, technical covariates, and first 10 genetic PCs, overall mental health problems were not significantly associated with either gut microbiome richness or the two diversity indices in children (Table 2). Most associations were negative in direction, meaning less diversity is found in the case of more mental health problems. Follow-up analyses examining specific mental health problems showed comparable findings.

### Child mental health and gut microbiome profiles

We analyzed single taxonomies associated with child mental health using univariate ANCOM-BC differential abundance models. We identified a total of 7 genera associated with either overall or specific mental health problems in adjusted models, based on *p* < 0.01 (Supplementary Table 3). These associations, however, did not survive correction for multiple testing.

A multivariate approach was performed to investigate if overall gut microbiome compositions are associated with mental health problems in children. A PERMANOVA test was performed on the pair-wise dissimilarities between samples to detect clustering of microbiome profiles associated with overall and specific phenotypes, while adjusting for covariates. No significant associations could be detected after multiple testing correction (Supplementary Table 4).

When we assessed all genera that were nominally significant in the ANCOM-BC univariate analysis (i.e., *Muribaculaceae unknown genus, Erysipelatoclostridium, Anaeroplasma, Tyzzerella, Eubacterium ruminantium group, Hungatella, Anaerotruncus*) (Fig. 1) for their effect on all overall and specific mental health problems in a heatmap, we observed the same direction of effect across all phenotypes. Furthermore, results from the multivariate analysis (PERMANOVA) was largely consistent with the respective univariate analyses, see overlap in Figure 1.

**Figure 1.**
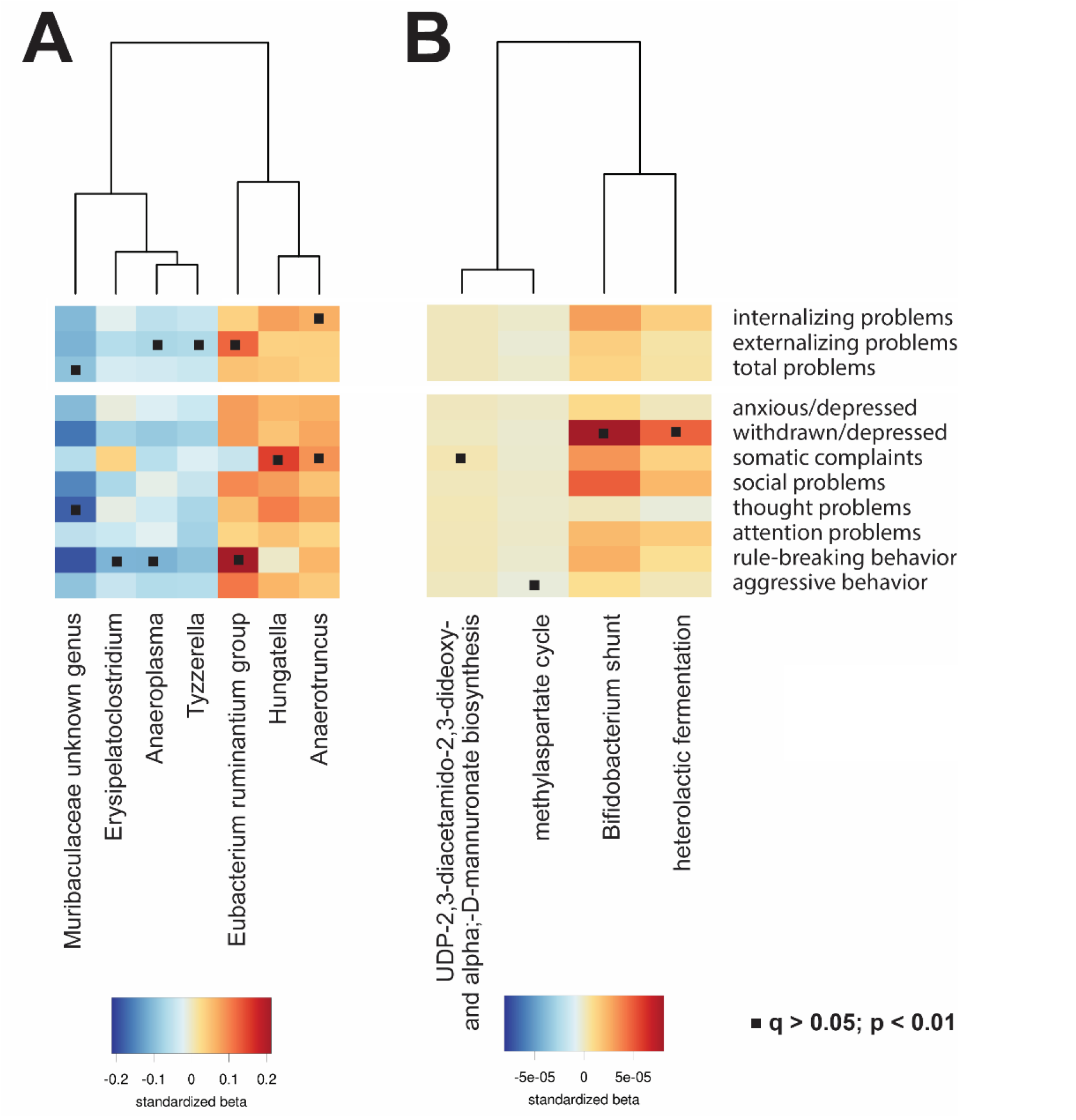
Associations of gut microbiome and microbial functions with mental health phenotypes. (A) Genus-level taxonomies that were nominally significant (p < 0.01 and q > 0.05; indicated by a square) associated with one of the mental health phenotypes are listed in the heatmap, after covariate adjustment. Color represents the standardized beta from the ANCOM-BC function. (B) Predicted microbial functions organized in MetCyc pathways that were nominally significant (p < 0.01 and q > 0.05; indicated by a dot) significant associated with one of the mental health phenotypes are listed in the heatmap, after covariate adjustment. Colors represent the beta obtained from the LM linear model function.

### Child mental health and gut microbial functions

The functional content of gut microbiome was predicted from the ASV-level abundance tables using the PICRUSt2 tool and combined in annotated MetaCyc pathways (Supplementary Table 5), while adjusting for covariates. No findings emerged for overall mental health problems symptoms. In follow-up analyses, nominally significant associations were observed for the withdrawn/depressed phenotype with two fermentation pathways that lead to the production of lactate (Bifidobacterium shunt and heterolactic fermentation). Furthermore, a nominally significant association was observed for methylasperate cycle with aggressive behavior; and for UDP-2,3-diacetamido-2,3-dideoxy-&alpha;-D-mannuronate biosynthesis with somatic complaints. (Fig. 1). As with other findings, these associations did not survive multiple testing correction.

## DISCUSSION

We examined associations between gut microbiome and child mental health problems at different taxonomic levels (alpha diversity measures, genus-level, and functional pathways), drawing on data from a large population-based study of almost 1,800 10-year-old children. Overall, we found no clear evidence of a cross-sectional association between gut microbiome composition and either overall or specific child mental health problems. The direction and magnitude of associations for each taxon was similar across mental health outcomes, reflecting the correlated nature of emotional and behavioral problems. Furthermore, although some of the identified associations were consistent with previous findings, implicating genera such as *Anaerotruncus, Anaeroplasma*, and *Hungatella*, none of these associations survived multiple-testing correction. We also did not identify any enriched microbial functional pathways in relation to child mental health problems. Together, our findings do not support a strong link between the gut microbiome and mental health in the general pediatric population.

We first examined the relationship between indices of gut microbial diversity and child mental health problems. We found that although both overall (e.g., externalizing problems) as well as specific (e.g., rule-breaking behavior) mental health problems were generally associated with lower gut microbial diversity and richness, these associations were not significant. While reduced microbial diversity has been robustly associated with variables such as higher age, BMI and antibiotic use, the relationship with mental health problems other than adult depression and autism spectrum disorder has been inconsistent (6, 7, 10). Our findings are in line with other population-based studies on the association between gut microbiome and mental health, which reported weak associations with diversity indices in children and adults (8, 9, 11). As a second step, we found that no single genus was associated with overall or specific mental health problems after multiple-testing correction. Associations also seemed to be non-specific, with effects being of similar magnitude and direction across outcomes. This is perhaps unsurprising given the known co-occurrence between mental health symptoms and highlights the importance of assessing multiple psychiatric outcomes concurrently. Such an approach, however, contrasts with what is most commonly done in the field, where individual psychiatric outcomes are typically examined in isolation (10). Indeed, a study reported associations with gut microbiome and having elevated symptoms in at least one domain of overall mental health problems (11), although associations with specific domains were not examined, thereby precluding comparisons across phenotypes. Like the single taxon or genus analyses, results from the multivariate PERMANOVA were not significant after multiple-testing correction, although taxa that best explained the mental health problems showed overlap with genera that were nominally significantly associated in the single taxon analyses.

Although findings did not survive multiple-testing correction, we highlight three genera that were most consistently associated with child mental health problems across analyses and that have been associated with psychiatric risk in previous literature. First, increased internalizing problems and somatic complaints were associated with higher abundance of the *Anaerotruncus* genus, in accordance with broader literature. Although only consisting of small individual studies, increased *Anaerotruncus* has been implicated in several psychiatric diagnoses, such as schizophrenia and anorexia nervosa (34), but also nominally in major depressive disorder (35). Further, we found that child externalizing problems, specifically rule-breaking behavior, associated with lower abundance of *Anaeroplasma*. Lower abundance of this genus has also been reported by animal experimental studies in relation to relevant phenotypes such as more dominant and less submissive behavior (36), as well as compulsive-like behaviors (37) and depression (38). However, these associations have thus far not been reported in other human samples. Finally, we found associations with increased abundances of *Hungatella* in relation to higher somatic complaints, a feature of internalizing problems. Interestingly, increased abundance of *Hungatella* has been previously found to associate with infant temperament (39), and notably also to depression in a large population-based sample of adults (9). Together, these findings point to *Hungatella* as an interesting candidate for future research, particularly in relation to features of depression and its developmental precursors in childhood.

As a final step, we performed a functional analysis to examine whether specific microbial functions may be enriched in relation to child mental health problems. We did not find any functional pathways to be enriched after multiple-testing correction for overall mental health problems. As for specific mental health problems, we do note that a nominally significant association was identified between aggressive behavior and the methylaspartate cycle. Multiple lines of evidence support a role of the N-methyl-D-aspartate receptor in the pathophysiology of several psychiatric disorders such as schizophrenia (40) and bipolar disorder (41), although exact mechanisms still remain unclear. Further, a nominally significant association emerged between withdrawn/depressed behavior and enrichment for two pathways independently involved in glucose fermentation (i.e., Bifidobacterium shunt and heterolactic fermentation). Glucose fermentation is linked to the production of lactate, and increased lactate is linked to energy metabolic dysfunction, which may be a key component to depressive symptoms (42). Supporting this is also evidence of increased lactate in patients experiencing a major depressive episode compared to healthy controls (43). While these findings are intriguing and link child mental health problems to metabolic markers and amino acids at the level of the gut microbiome, it is not known which specific enzymes in the pathway account for this enrichment. As such, future studies will be needed to test the robustness or even the existence of these associations and their potential functional relevance.

Overall, our findings point to no clear links between the gut microbiome and mental health problems in children. This contrasts with the existing literature, which has been rife with reports of significant associations – although few consistent findings have emerged based on recent systematic reviews (6, 7, 10). Weak to no associations may be explained by several factors. First, we examined mental health symptoms dimensionally in the general population, as opposed to most previous research based on case-control comparisons in patient samples. Dimensional analyses of continuous variables have the advantage of modelling the full spectrum of symptoms as a continuum in the general population. However, if associations with the gut microbiome manifest only at more severe ends of the symptom spectrum, our study may have lacked the symptom severity necessary to detect such associations. Second, we investigated common mental health problems in children, which precede the peak onset of several psychiatric disorders that have been linked to the gut microbiome in adults, such as major depression or schizophrenia. Here, we do not find significant evidence that gut microbiome associates with developmental precursors for these disorders (e.g., emotional and thought problems) in childhood. On the one hand, it is possible that associations with the gut microbiome only emerge during specific developmental periods, for instance once a psychiatric disorder is fully manifested. On the other hand, it is also possible that differences in composition observed in adults may be more likely a consequence rather than a risk factor for these psychiatric disorders, for example because of dietary and lifestyle changes associated with the disorder, such as what has been seen in autism spectrum disorder (44). Finally, a possible interpretation of our results is that the gut microbiome does not substantially affect mental health symptoms, and that previously reported associations may have been biased by factors such as unmeasured confounding, small sample sizes and inadequate adjustment for false positives. Yet, at a broader level, the small effect sizes observed in our study are in line with what has been reported by other population *omics* research on mental health, including at the genetic, epigenetic, and transcriptomic level (6, 7, 10).

### Limitations and future directions

Our findings should be interpreted considering several limitations. First, we focused on common mental health problems in children and as such did not assess autism spectrum disorder (ASD), which is currently the most examined child brain-based disorder in relation to the gut microbiome (6). In future, it will be important to test whether an association between gut microbiome composition and ASD symptoms can be identified in the general pediatric population, but associations may also be tested for other brain-related outcomes. Second, our study, like many others in the field, was cross-sectional. An important step for future research will be the assessment of longitudinal data at repeated time points, to clarify the direction of associations between the gut microbiome and mental health, and to test whether associations emerge during specific developmental periods. Of note, one study that did assess gut microbiome repeatedly (at age 1, 6 and 12 months) and associated this to mental health at 2 years established only proximal associations (i.e., with gut microbiome at age 12 months, and not for microbiome assessed at age 1 and 6 months) (11). Third, we were not able to adjust for potentially important covariates such as medication use, although participants were excluded if they recently were administered antibiotics and use of psychiatric medication in this pediatric population is expected to be low compared to patient samples. In addition, we do not correct for dietary patterns, which may lead to less precise estimates. Fourth, gut microbiome was estimated from stool samples; however, we do not know how well stool samples reflect the full gastrointestinal tract microbiome. Fifth, although we adjusted for duration of exposure as a covariate, we collected stool samples at room temperature, which can affect survival of anaerobes (15). Finally, we could not meta-analyze our results with other studies due to a lack of available population-based pediatric cohorts with gut microbiome and mental health data in middle childhood. It is thus possible that associations between the gut microbiome and child mental health are too subtle to be detected with our current sample alone. However, with data on almost 1,800 children, our study is substantially larger than published research (6, 7, 10), making an important contribution to existing literature.

## Conclusion

Based on this large, population-based study, we find little evidence of an association between the gut microbiome and common mental health problems in children. Our study does not definitively refute a link between the gut microbiome and child mental health symptoms but indicates that associations are likely of small magnitude in the general pediatric population. In future, larger studies may be necessary to detect subtle associations with greater statistical power. Furthermore, the use of longitudinal data from early life to adulthood will mark a crucial step for understanding the role of the gut microbiome in the development of psychiatric symptoms and characterizing how associations between these factors unfold over time.

## Funding

The Generation R Study is conducted by the Erasmus MC in close collaboration with the Erasmus University Rotterdam, the Municipal Health Service Rotterdam area, Rotterdam, and the Rotterdam Homecare Foundation, Rotterdam. We acknowledge the contribution of children and parents, general practitioners, hospitals, midwives, and pharmacies in Rotterdam. The general design of Generation R Study was made possible by financial support from the Erasmus MC, Rotterdam, the Erasmus University Rotterdam, the Netherlands Organization for Health Research and Development (ZonMW), the Netherlands Organization for Scientific Research (NWO), the Ministry of Health, Welfare and Sport and the Ministry of Youth and Families. This project also received funding from the European Union’s Horizon 2020 research and innovation programme under the following grant agreements: (No 848158 [EarlyCause]; No. 733206 [LIFECYCLE]). CC is supported by the European Research Council (ERC) under the European Union’s Horizon 2020 Research and Innovation Programme [grant agreement No 101039672 [TEMPO]). Djawad Radjabzadeh was funded by an Erasmus MC mRACE grant “Profiling of the human gut microbiome”. The authors declare no conflicts of interest.

## Acknowledgements

The generation and management of the 16S microbiome data for the Generation R Study was executed by the Human Genotyping Facility of the Genetic Laboratory of the Department of Internal Medicine, Erasmus MC, University Medical Center Rotterdam, the Netherlands. We thank Nahid El Faquir and Jolande Verkroost-Van Heemst for their help in sample collection and registration and Kamal Arabe, Hedayat Razawy, Karan Singh Asra, Pelle van der Wal, Sergio Chavez and Djawad Radjabzadeh for their help in DNA isolation and sequencing, and Joost Verlouw, Dr. Constanza Vallerga and Marijn Verkerk for their help with the bioinformatic analyses. We thank Ruolin Li, Dr. Cindy Boer, Dr. Robert Kraaij, Dr. Carolina Medina-Gomez and Prof. Joyce van Meurs for overseeing the quality control of the generated datasets. Furthermore, we express our gratitude to Drs. Jeroen Raes and Jun Wang (Katholieke Universiteit Leuven, Belgium) for their guidance in 16S rRNA profiling and dataset generation.

Supplementary information is available at Molecular Psychiatry’s website.

## Notes

### Competing Interest Statement

The authors have declared no competing interest.

