## Supplementary Table for "The gut microbiome and child mental health: a population-based study"

**SUPPLEMENTARY METHODS**

**Stool microbiome data**

Stool samples were collected at a mean age of 9.8 years (SD = 0.3) and DNA isolation, 16S rRNA profiling and filtering were performed as described (1). In brief, samples were collected at home by the participants and an aliquot of approximately 1 gram was sent to the Erasmus MC research center by regular mail. Samples were stored at -20°C before DNA was isolated using the automated Arrow Stool DNA isolation kit (Isogen Life Science, De Meern, The Netherlands) after bead beating with 0.1 mm silica beads (MP Biomedicals, LLC, Bio Connect Life Sciences BV, Huissen, The Netherlands). The V3 and V4 variable regions of the 16S rRNA gene were amplified using the 309F-806R primer pair and dual indexing (2) followed by Illumina MiSeq sequencing (Illumina Inc., San Diego, CA) on the V3 flowcell (MiSeq Reagent Kit v3, 2 x 300 bp) at an average depth of 50,000 read-pairs per sample.

**Data processing**

Raw reads from Illumina MiSeq were demultiplexed using a custom script to separate sample fastq files based on the dual index. Primers, barcodes and heterogeneity spacers were trimmed off using tagcleaner v0.16 (3). Trimmed fastq files were loaded into R v4.0.0 (4) with the DADA2 (5) package. Quality filtering was performed in DADA2 using the following criteria: trim=0, maxEE=c(2,2), truncQ=2, rm.phix=TRUE. Filtered reads were run through the DADA2 Amplicon Sequence Variant (ASV) assignment tool to denoise, cluster and merge the reads. ASVs were assigned a taxonomy from the SILVA version 138.1 rRNA database (6) using the RDP naïve Bayesian classifier (7). The resulting data tables were combined into a phyloseq object using Phyloseq (8) and a phylogenetic tree was generated using the Phangorn R package (9) based on the sequences of the ASVs and added to the phyloseq object. PICRUSt2 prediction of functional MetaCyc microbial pathways was performed in R using PICRUSt2 (10) with the default EPA-NG (11) placement option and MinPath (12) biological pathway reconstruction based on Enzyme Commission (EC) numbers (13).

**Web resources**

Generation R study: https://www.generationr.nl/researchers/; QIIME: http://qiime2.org/; ; TAGcleaner: http://tagcleaner.sourceforge.net/; Vegan R package: https://cran.r-project.org/web/packages/vegan; PICRUSt2: https://huttenhower.sph.harvard.edu /picrust/

**Supplemental Table 1. Sample filtering**

| Filter | *N* |
| --- | --- |
| None (wave at age 9 participants) | 2,526 |
| Technical covariates available | 2,275 |
| Not exceeding time in mail (days) > 5 | 2,187 |
| Child mental health phenotype available | 1,948 |
| Genetic PCs / consent available | 1,784 |

PCs: principal components

**Supplemental Table 2. Microbiome characteristics**

|  | | % | mean (SD) | *N* |
| --- | --- | --- | --- | --- |
| Season of sample production | |  |  |  |
|  | Spring (%) | 34.8 |  |  |
|  | Summer (%) | 21.5 |  |  |
|  | Autumn (%) | 22.9 |  |  |
|  | Winter (%) | 20.9 |  |  |
| Time in the mail | |  |  |  |
|  | 1 day (%) | 24.7 |  |  |
|  | 2 days (%) | 35.5 |  |  |
|  | 3 days (%) | 18.1 |  |  |
|  | 4 days (%) | 15.9 |  |  |
|  | 5 days (%) | 5.8 |  |  |
| DNA isolation batch | |  |  |  |
|  | Batch 0 (%) | 91.4 |  |  |
|  | Batch 1 (%) | 8.6 |  |  |
| Sequencing batch | |  |  |  |
|  | Batch 0 (%) | 84.4 |  |  |
|  | Batch 1 (%) | 0.2 |  |  |
|  | Batch 2 (%) | 3.4 |  |  |
|  | Batch 3 (%) | 12 |  |  |
| Number of reads | |  | 20,196 (11,918) |  |
| Alpha diversities | |  |  |  |
|  | observed ASVs |  | 149.10 (49.2) |  |
|  | Shannon |  | 3.98 (0.42) |  |
|  | Inversed Simpson |  | 31.50 (14.18) |  |
| Number of taxa | |  |  |  |
|  | ASVs |  | 149.1 (49.2) | 1,578 |
|  | Species |  | 84.2 (20.1) | 305 |
|  | Genus |  | 61.1 (14.6) | 188 |
|  | Phylum |  | 5.2 (1.1) | 10 |

ASVs: amplicon sequence variants

**Supplemental Table 3. Single genus differential abundance analysis of child mental health phenotypes: ANCOM-BC differential abundance analyses**

| Child mental health | Genus |  | B | se | *p* | fdr |
| --- | --- | --- | --- | --- | --- | --- |
| Anxious/depressed | Anaerotruncus_ASV850 | | 0.06 | 0.027 | .017 | .904 |
| Withdrawn/depressed | NA_ASV109 | | -0.16 | 0.069 | .018 | .747 |
|  | Anaeroplasma_ASV334 | | -0.09 | 0.038 | .022 | .747 |
|  | Blautia_ASV15 | | -0.07 | 0.033 | .027 | .747 |
|  | Anaerotruncus_ASV850 | | 0.07 | 0.034 | .028 | .747 |
|  | Colidextribacter_ASV323 | | -0.10 | 0.049 | .043 | .780 |
| Somatic complaints | Hungatella_ASV266 | | 0.15 | 0.054 | .005 | .363 |
|  | Anaerotruncus_ASV850 | | 0.09 | 0.033 | .007 | .363 |
|  | Coprobacillus_ASV661 | | 0.07 | 0.032 | .024 | .644 |
|  | DTU089_ASV461 | | 0.08 | 0.038 | .026 | .644 |
|  | NA_ASV225 | | 0.10 | 0.045 | .030 | .644 |
| Social problems | NA_ASV109 | | -0.15 | 0.061 | .015 | .970 |
|  | Erysipelatoclostridium_ASV909 | | -0.07 | 0.035 | .047 | .970 |
|  | Anaerotruncus_ASV850 | | 0.05 | 0.030 | .096 | .970 |
|  | Lachnospiraceae_AC2044_group_ASV226 | | 0.07 | 0.045 | .099 | .970 |
|  | Hungatella_ASV266 | | 0.08 | 0.050 | .103 | .970 |
| Thought problems | NA_ASV109 | | -0.18 | 0.066 | .006 | .466 |
|  | Anaerotruncus_ASV850 | | 0.08 | 0.031 | .011 | .466 |
|  | Desulfovibrio_ASV420 | | 0.10 | 0.042 | .013 | .466 |
|  | Erysipelotrichaceae_UCG_003_ASV120 | | -0.11 | 0.047 | .025 | .563 |
|  | Lachnospiraceae_ND3007_group_ASV100 | | -0.10 | 0.048 | .031 | .563 |
|  | Hungatella_ASV266 | | 0.11 | 0.049 | .032 | .563 |
|  | Tyzzerella_ASV414 | | -0.07 | 0.035 | .041 | .625 |
| Attention problems | Tyzzerella_ASV414 | | -0.08 | 0.032 | .018 | .986 |
|  | Eubacterium_eligens_group_ASV50 | | 0.10 | 0.047 | .032 | .986 |
|  | Veillonella_ASV454 | | 0.08 | 0.039 | .042 | .986 |
| Rule breaking behavior | Anaeroplasma_ASV334 | | -0.11 | 0.040 | .006 | .283 |
|  | Eubacterium_ruminantium_group_ASV67 | | 0.21 | 0.078 | .007 | .283 |
|  | Erysipelatoclostridium_ASV909 | | -0.11 | 0.041 | .008 | .283 |
|  | NA_ASV109 | | -0.19 | 0.076 | .013 | .338 |
|  | Ruminococcus_ASV41 | | 0.12 | 0.050 | .019 | .357 |
|  | Tyzzerella_ASV414 | | -0.09 | 0.041 | .022 | .358 |
|  | Romboutsia_ASV21 | | -0.14 | 0.064 | .027 | .358 |
|  | Prevotella_9_ASV23 | | 0.26 | 0.121 | .029 | .358 |
|  | Blautia_ASV15 | | -0.07 | 0.033 | .030 | .358 |
| Aggressive behavior | NA_ASV225 | | 0.09 | 0.033 | .010 | .708 |
|  | Anaeroplasma_ASV334 | | -0.06 | 0.027 | .020 | .708 |
|  | Tyzzerella_ASV414 | | -0.06 | 0.027 | .025 | .708 |
|  | Eubacterium_ruminantium_group_ASV67 | | 0.11 | 0.051 | .033 | .708 |
|  | NA_ASV440 | | -0.08 | 0.039 | .033 | .708 |
| Internalizing problems | Anaerotruncus_ASV850 | | 0.07 | 0.023 | .004 | .371 |
|  | Coprobacillus_ASV661 | | 0.05 | 0.023 | .026 | .533 |
|  | NA_ASV109 | | -0.10 | 0.048 | .029 | .533 |
|  | Colidextribacter_ASV323 | | -0.07 | 0.033 | .029 | .533 |
|  | Prevotella_9_ASV23 | | 0.16 | 0.077 | .033 | .533 |
|  | Anaeroplasma_ASV334 | | -0.05 | 0.025 | .034 | .533 |
|  | Hungatella_ASV266 | | 0.08 | 0.037 | .035 | .533 |
| Externalizing problems | Anaeroplasma_ASV334 | | -0.07 | 0.025 | .005 | .323 |
|  | Tyzzerella_ASV414 | | -0.07 | 0.026 | .009 | .323 |
|  | Eubacterium_ruminantium_group_ASV67 | | 0.12 | 0.047 | .009 | .323 |
|  | Erysipelatoclostridium_ASV909 | | -0.06 | 0.025 | .014 | .351 |
|  | NA_ASV109 | | -0.11 | 0.046 | .017 | .357 |
|  | Romboutsia_ASV21 | | -0.09 | 0.039 | .028 | .503 |
|  | NA_ASV225 | | 0.06 | 0.031 | .043 | .652 |
| Total problems | NA_ASV109 | | -0.09 | 0.032 | .003 | .356 |
|  | Anaerotruncus_ASV850 | | 0.04 | 0.015 | .013 | .557 |
|  | Tyzzerella_ASV414 | | -0.04 | 0.018 | .016 | .557 |
|  | Romboutsia_ASV21 | | -0.06 | 0.027 | .025 | .654 |
|  | Anaeroplasma_ASV334 | | -0.04 | 0.017 | .033 | .677 |
|  | Lachnospiraceae_ND3007_group_ASV100 | | -0.05 | 0.024 | .039 | .687 |
|  | Prevotella_9_ASV23 | | 0.10 | 0.051 | .047 | .713 |

Genus-level taxonomies that were borderline (p < 0.05) significant associated with the mental health phenotypes. ANCOM-BC linear regression models with child mental health phenotypes and genus abundance. Model: sqrt(child mental health phenotype) ~ genus + age + sex + BMI + self-reported antibiotics use + maternal education + time in mail + season of stool production + DNA isolation batch + sequencing run batch + number of sequencing reads + first 10 genetic PCs. Values are pooled from 30 imputed datasets. Sqrt: square root transformed; PCs: principal components; se: standard error; fdr: false discovery rate corrected *p*-value using the Benjamini-Hochberg correction.

**Table 4. Associations between child mental health phenotypes and overall gut microbiome composition: PERMANOVA**

| Child mental health | *F* | *p* | RIV |
| --- | --- | --- | --- |
| Empirical scales |  |  |  |
| Anxious/depressed | 1.020 | .407 | 0.055% |
| Withdrawn/depressed | 0.934 | .566 | 0.051% |
| Somatic complaints | 1.202 | .179 | 0.066% |
| Social problems | 0.866 | .689 | 0.047% |
| Thought problems | 0.974 | .487 | 0.053% |
| Attention problems | 0.957 | .519 | 0.052% |
| Rule-breaking behavior | 1.370 | .073 | 0.074% |
| Aggressive behavior | 0.874 | .675 | 0.047% |
| Broadband and total scales |  |  |  |
| Internalizing problems | 1.197 | .181 | 0.065% |
| Externalizing problems | 1.124 | .260 | 0.061% |
| Total problems | 1.184 | .193 | 0.064% |

PERMANOVA of child mental health phenotypes and gut microbiome composition. Model: distance matrix ~ age + sex + BMI + self-reported antibiotics use + maternal education + time in mail + season of stool production + DNA isolation batch + sequencing run batch + number of sequencing reads + first 10 genetic PCs + sqrt(child mental health phenotype). The distance matrix was generated from the filtered genus-level taxonomy table after centered log ratio transformation. Values are medians of 30 iterations that include different imputed datasets. Sqrt: square root; PCs: principal components.

**Table 5. Single pathway differential abundance analysis of child mental health phenotypes**

| Child mental health | Pathway | MetaCyc pathway code | B | se | *p* | fdr |
| --- | --- | --- | --- | --- | --- | --- |
| Anxious/depressed | ethylmalonyl-CoA pathway | PWY.5741 | 2.69E-06 | 1.19E-06 | .024 | .998 |
|  | L-leucine degradation I | LEU.DEG2.PWY | 2.72E-06 | 1.21E-06 | .025 | .998 |
|  | superpathway of N-acetylglucosamine. N-acetylmannosamine and N-acetylneuraminate degradation | GLCMANNANAUT.PWY | -5.02E-05 | 2.32E-05 | .031 | .998 |
|  | superpathway of L-aspartate and L-asparagine biosynthesis | ASPASN.PWY | 3.45E-05 | 1.71E-05 | .044 | .998 |
|  | purine ribonucleosides degradation | PWY0.1296 | -5.78E-05 | 2.87E-05 | .044 | .998 |
| Somatic complaints | UDP-2.3-diacetamido-2.3-dideoxy-&alpha;-D-mannuronate biosynthesis | PWY.7090 | 2.72E-06 | 9.15E-07 | .003 | .920 |
|  | photorespiration | PWY.181 | 2.77E-06 | 1.20E-06 | .021 | .999 |
|  | chitin derivatives degradation | PWY.6906 | 2.64E-06 | 1.28E-06 | .040 | .999 |
| Withdrawn/depressed | P124.PWY | Bifidobacterium shunt | 8.10E-05 | 3.09E-05 | .009 | .990 |
|  | P122.PWY | heterolactic fermentation | 4.77E-05 | 1.83E-05 | .009 | .990 |
|  | PWY.5005 | biotin biosynthesis II | -1.64E-05 | 8.27E-06 | .047 | .990 |
| Social problems | superpathway of &beta;-D-glucuronide and D-glucuronate degradation | GLUCUROCAT.PWY | 4.69E-05 | 1.83E-05 | .011 | .889 |
|  | thiazole biosynthesis II (Bacillus) | PWY.6891 | 3.91E-05 | 1.71E-05 | .023 | .889 |
|  | superpathway of hexuronide and hexuronate degradation | GALACT.GLUCUROCAT.PWY | 3.56E-05 | 1.58E-05 | .024 | .889 |
|  | chitin derivatives degradation | PWY.6906 | 2.73E-06 | 1.21E-06 | .025 | .889 |
|  | glycolysis I (from glucose 6-phosphate) | GLYCOLYSIS | -5.40E-05 | 2.58E-05 | .037 | .889 |
| Thought problems | UDP-2.3-diacetamido-2.3-dideoxy-&alpha;-D-mannuronate biosynthesis | PWY.7090 | 2.11E-06 | 8.79E-07 | .017 | .982 |
|  | glycogen degradation I (bacterial) | GLYCOCAT.PWY | -6.19E-05 | 2.96E-05 | .037 | .982 |
|  | superpathway of UDP-N-acetylglucosamine-derived O-antigen building blocks biosynthesis | PWY.7332 | 1.83E-05 | 9.02E-06 | .043 | .982 |
| Attention problems | UDP-2.3-diacetamido-2.3-dideoxy-&alpha;-D-mannuronate biosynthesis | PWY.7090 | 1.78E-06 | 7.68E-07 | .021 | .842 |
|  | dTDP-L-rhamnose biosynthesis I | DTDPRHAMSYN.PWY | -4.56E-05 | 2.06E-05 | .027 | .842 |
|  | peptidoglycan biosynthesis IV (Enterococcus faecium) | PWY.6471 | -5.83E-05 | 2.78E-05 | .036 | .842 |
|  | superpathway of arginine and polyamine biosynthesis | ARG.POLYAMINE.SYN | 4.11E-05 | 2.02E-05 | .042 | .842 |
|  | nitrate reduction VI (assimilatory) | PWY490.3 | -3.47E-05 | 1.72E-05 | .044 | .842 |
|  | starch degradation V | PWY.6737 | -5.11E-05 | 2.54E-05 | .044 | .842 |
| Rule-breaking behavior | photorespiration | PWY.181 | 3.63E-06 | 1.46E-06 | .013 | .955 |
|  | superpathway of sulfur oxidation (Acidianus ambivalens) | PWY.5304 | -7.12E-05 | 2.89E-05 | .014 | .955 |
|  | tetrapyrrole biosynthesis II (from glycine) | PWY.5189 | -6.19E-05 | 2.67E-05 | .020 | .955 |
|  | GDP-D-glycero-&alpha;-D-manno-heptose biosynthesis | PWY.6478 | 3.59E-05 | 1.63E-05 | .028 | .955 |
|  | sucrose degradation IV (sucrose phosphorylase) | PWY.5384 | -7.52E-05 | 3.48E-05 | .031 | .955 |
|  | purine ribonucleosides degradation | PWY0.1296 | -7.82E-05 | 3.72E-05 | .036 | .955 |
|  | GDP-mannose biosynthesis | PWY.5659 | 5.32E-05 | 2.55E-05 | .037 | .955 |
|  | glycerol degradation to butanol | PWY.7003 | -2.79E-05 | 1.35E-05 | .039 | .955 |
|  | tetrapyrrole biosynthesis I (from glutamate) | PWY.5188 | -5.78E-05 | 2.86E-05 | .043 | .955 |
|  | UDP-2.3-diacetamido-2.3-dideoxy-&alpha;-D-mannuronate biosynthesis | PWY.7090 | 2.05E-06 | 1.03E-06 | .048 | .955 |
| Aggressive behavior | methylaspartate cycle | PWY.6728 | -3.81E-06 | 1.24E-06 | .002 | .681 |
|  | chlorophyllide a biosynthesis II (anaerobic) | PWY.5531 | -4.46E-07 | 2.21E-07 | .044 | .892 |
|  | chlorophyllide a biosynthesis III (aerobic. light independent) | PWY.7159 | -4.46E-07 | 2.21E-07 | .044 | .892 |
| Internalizing problems | biotin biosynthesis II | PWY.5005 | -1.28E-05 | 5.71E-06 | .026 | .997 |
| Externalizing problems | methylaspartate cycle | PWY.6728 | -2.83E-06 | 1.14E-06 | .013 | .893 |
|  | nitrate reduction VI (assimilatory) | PWY490.3 | -3.29E-05 | 1.41E-05 | .019 | .893 |
|  | hexitol fermentation to lactate. formate. ethanol and acetate | P461.PWY | -3.43E-05 | 1.56E-05 | .028 | .893 |
|  | peptidoglycan biosynthesis IV (Enterococcus faecium) | PWY.6471 | -4.91E-05 | 2.28E-05 | .031 | .893 |
| Total problems | peptidoglycan biosynthesis IV (Enterococcus faecium) | PWY.6471 | -3.67E-05 | 1.58E-05 | .020 | .990 |
|  | methylaspartate cycle | PWY.6728 | -1.65E-06 | 7.91E-07 | .038 | .990 |
|  | nitrate reduction VI (assimilatory) | PWY490.3 | -1.98E-05 | 9.76E-06 | .042 | .990 |
|  | myo-inositol degradation I | P562.PWY | -4.72E-06 | 2.33E-06 | .043 | .990 |
|  | UDP-2.3-diacetamido-2.3-dideoxy-&alpha;-D-mannuronate biosynthesis | PWY.7090 | 8.79E-07 | 4.37E-07 | .044 | .990 |
|  | arginine. ornithine and proline interconversion | ARGORNPROST.PWY | -4.89E-06 | 2.48E-06 | .049 | .990 |

Linear modeling of child mental health phenotype and single pathways. Model: relative pathway abundance ~ sqrt(child mental health phenotype) + age + sex + BMI + self-reported antibiotics use + maternal education + time in mail + season of stool production + DNA isolation batch + sequencing run batch + number of sequencing reads + first 10 genetic PCs. Values are pooled over 30 iterations that include different imputed datasets. Sqrt: square root; PCs: principal components; se: standard error; fdr: false discovery rate corrected *p*-value using the Benjamini-Hochberg correction.
